# Cationic polymethacrylate-copolymer acts as an agonist for β-amyloid and antagonist for amylin fibrillation

**DOI:** 10.1101/401687

**Authors:** Bikash R. Sahoo, Takuya Genjo, Andrea K. Stoddard, Kazuma Yasuhara, Carol A. Fierke, Ayyalusamy Ramamoorthy

**Affiliations:** Biophysics and Department of Chemistry, University of Michigan, Ann Arbor, MI 48109, USA; Graduate School of Materials Science, Nara Institute of Science and Technology, Nara 6300192, Japan; Department of Chemistry, Texas A&M University, College Station, TX 77843, USA

**Keywords:** Amyloid-beta, IAPP, Alzheimer’s disease, Type-II diabetes mellitus, Protein misfolding

## Abstract

In human, amyloid-beta (Aβ) and islet amyloid polypeptide (hIAPP) aggregations are linked to Alzheimer’s disease and Type-2 Diabetes, respectively. There is significant interest in better understanding the aggregation process by using chemical tools. Here, we show the ability of a cationic polymethacrylate-copolymer (PMAQA) to quickly induce β-hairpin structure and promote fibrillation in Aβ40, and to constrain the conformational plasticity of hIAPP for several days and inhibit its aggregation at sub-micromolar concentrations. NMR experiments and atomistic molecular dynamics simulations reveal that PMAQA electrostatically interacts with Aβ40’s Glu22 and Asp23 followed by β-sheet induction while it binds strongly to the closest proximity of amyloid core domain (NFGAIL) of hIAPP and restrain its structural rearrangement. This study provides a valuable approach to develop polymer-based anti-amyloid inhibitors that may diminish the population of intermediates of Aβ40 or hIAPP.

## Introduction

Self-assembly of amyloidogenic proteins is involved in numerous neurodegenerative diseases and Type-2 Diabetes (T2D).^1,2^ Nevertheless, our current understanding of the role of protein aggregation in the pathogenesis of such diseases remains elusive.^3^ Despite the recent advancements in high-throughput screening of several anti-amyloidogenic compounds,^4^ there is no treatment for the protein aggregation based disorders including Alzheimer’s disease (AD) and Type-2 Diabetes (T2D).^5^ Human amyloid-beta (Aβ) and islet amyloid polypeptide (hIAPP or amylin) aggregations are linked to AD and T2D, respectively.^6^ The conserved amyloidogenic nature of Aβ and hIAPP aggregation has been the subject of intense research to establish a pathological correlation^6^. The sequential protein aggregation mechanism from a water soluble monomer to insoluble amyloid fibers triggering AD and T2D still remains elusive. However, several studies observed a conserved pathway where both Aβ and hIAPP monomers aggregate to form pre-fibrillar toxic oligomers followed by matured β-sheet rich fiber structures.^7^ Thus, the sequential reaction products of Aβ or hIAPP aggregation pathways have been targeted to design potential inhibitors to interrupt amyloid formation. Moreover, substantial research effort has been devoted in developing strategies to reduce the formation of toxic intermediates of Aβ or hIAPP. In response to this, several amyloid inhibitors or modulators have been clinically tested for AD or T2D treatment.

However, small molecule inhibitors or modulators targeting amyloidosis have recently faced several clinical trial challenges.^[8,9]^ Among several compounds, the chemically conserved scaffold molecules characterized by multi aromatic groups have been tested *in vitro*^[10,11]^ or *in vivo*.^12^ These compounds have been reported to modulate the amyloid aggregation pathways. But the poor solubility and bioavailability affect their therapeutic nature and recently have been shown to be overcome using nanocarriers.^13–15^ Although, their mechanism of action remains unknown, hydrogen bonding, hydrophobic interactions and π-π stacking are thought to be the major driving forces for their inhibitory mechanism of action.^16,17^ Despite substantial efforts, no significant new drugs have been discovered against these most tenacious and unnerving medical disorders. The successive failures of small molecule compounds directs researchers to develop new anti-amyloidogenic molecules such as polymers, peptoids, nanoparticles, molecular chaperones etc.^18–20^ Among them, several polymers characterized by their ionic properties have been tested to investigate their activities on amyloidogenic aggregation. Notably, amine containing polymers, polyamino acids, cationic surfactants and cellular polyamines have been observed to modulate Aβ aggregation.^21–23^ Similarly, controlled aggregation kinetics and toxicity of hIAPP using star-polymers and polymer-nanodiscs have been studied recently^19,24^. Here we demonstrate the modulation of amyloid aggregation pathways for hIAPP and Aβ40 using a polymethacrylate (PMAQA) derived copolymer (Fig. S1) that has been implicated in several biological studies including lipid-nanodiscs formation, enhancement of drug-delivery, bioavailability and microencapsulation.^24–26^

## Results and Discussion

Far-UV circular dichroism (CD) spectra revealed a gradual structural transition (unfolded to folded) in Aβ40 titrated with PMAQA as indicated by a change in CD minima at ≈200 nm. At 1:5 PMAQA to Aβ40 molar ratio (Fig. 1A), a partial helical CD spectrum containing 9.4/26.1% of α/β secondary contents (as estimated by BESTSEL^27^) was observed. A further increase in PMAQA concentration to 1:1 PMAQA:Aβ40 molar ratio marginally increased the β-sheet contents in Aβ40 (7.7/28.2% of α/β content). The decrease in the CD minima at ≈200 nm with an increasing concentration of PMAQA indicated its activity on promoting Aβ40’s aggregation. Remarkably, unlike Aβ40, no significant conformational changes were observed in hIAPP when titrated with PMAQA even at two equivalent higher molar concentration of hIAPP (Fig. 1B). Thioflavin-T (ThT) based fluorescence aggregation assays of Aβ40 or hIAPP over 4 days were in line with the CD observation. As shown in Fig. 1(C and D), PMAQA promoted and inhibited Aβ40 and hIAPP aggregations, respectively. At the lowest concentration of PMAQA (1 µM), a significant difference in the lag-times of Aβ40 and hIAPP was ascertained (Fig. 1C and D, blue traces). At equimolar polymer:peptide concentration, Aβ40 aggregation was ≈6 times faster whereas hIAPP aggregation was significantly inhibited (Fig. 1C and D, pink traces).

**Fig 1.**
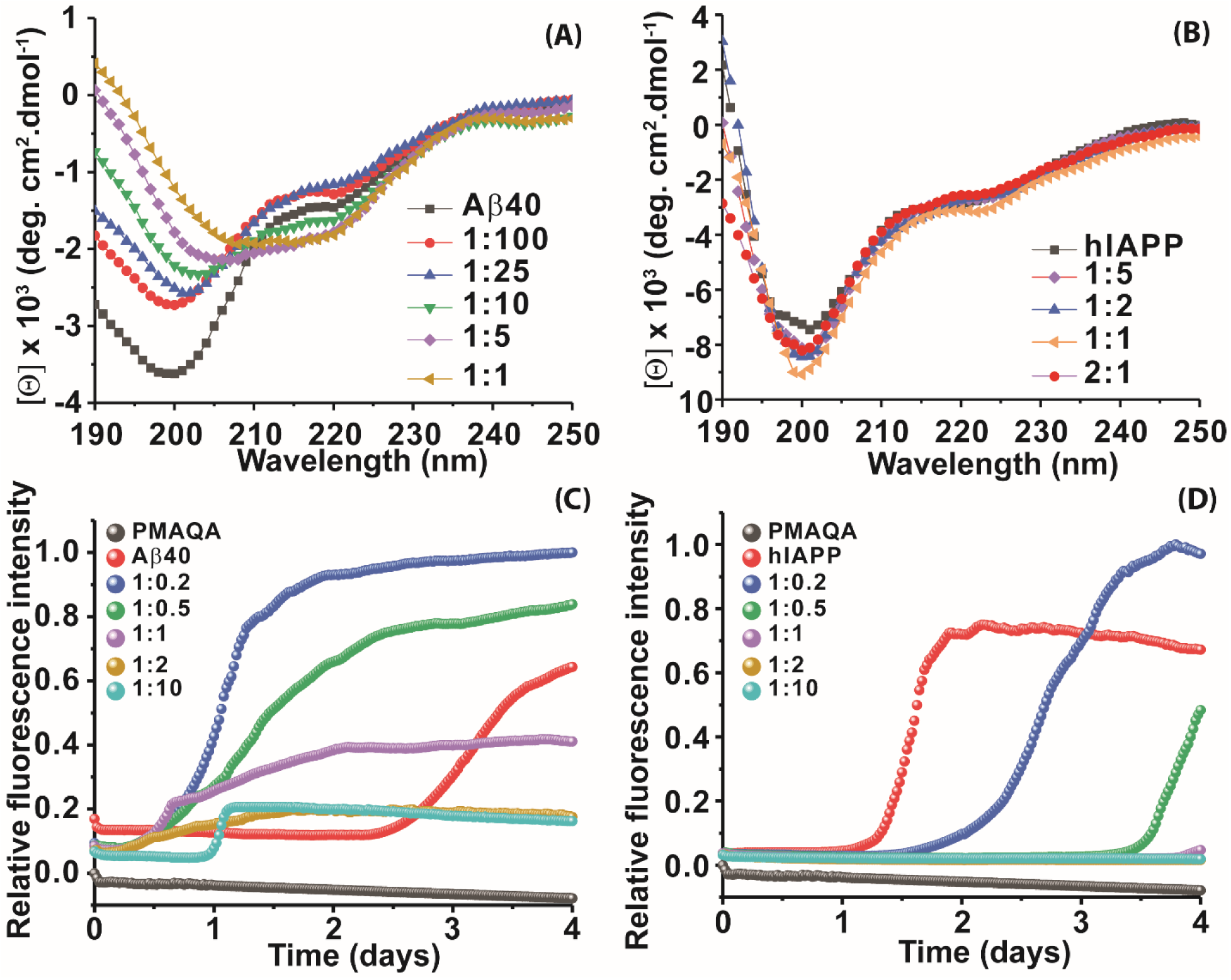
Effect of PMAQA on the conformation and aggregation of Aβ40 or hIAPP. (A-B) Far-UV CD analysis of 25 μM peptide (Aβ40 in 10 mM sodium phosphate buffer, pH 7.4 or hIAPP in 30 mM sodium acetate buffer, pH 5.4) in absence or presence of PMAQA at the indicated PMAQA to peptide molar concentration ratios. (C-D) Relative ThT fluorescence of 5 μM Aβ40 or 10 µM IAPP in presence of PMAQA at the indicated peptide to polymer molar ratios.

We next monitored the changes in the secondary structure of Aβ40 or hIAPP at 1:1.5 peptide:PMAQA molar ratio incubated at room temperature up to 5 days. The time-lapse CD spectra of Aβ40 dissolved in 10 mM sodium phosphate buffer, pH 7.4 or hIAPP dissolved in 30 mM sodium acetate buffer, pH 5.4, in absence of PMAQA showed a sequential structural transition from a random-coil (negative peak ≈200 nm) to β-structured fiber (Figs. 2C and S2A). While Aβ40 in solution (shown in Fig. S2A) are reported to exhibit a sequential structural change to form cross β-sheet structures over several days,^28^ interestingly, PMAQA showed rapid β-sheet induction within minutes when directly titrated with 1.5 equivalent molar concentration of PMAQA (Fig. 2A). The observed rapid structural change correlates well with fibrillation (Figs.1C and 2A) and indicates a reduction in the level of potential Aβ40 oligomers that are reported to be neurotoxic.^28,29^While discovery and design of small molecule Aβ40 inhibitors are of increasing interest, the observed role of PMAQA in converting misfolded Aβ40 rapidly to a stable β-sheet structure species could be useful to probe the mechanism of Aβ40 aggregation and the role of β-sheet in the formation of polymorphic fibers in AD.

**Fig 2.**
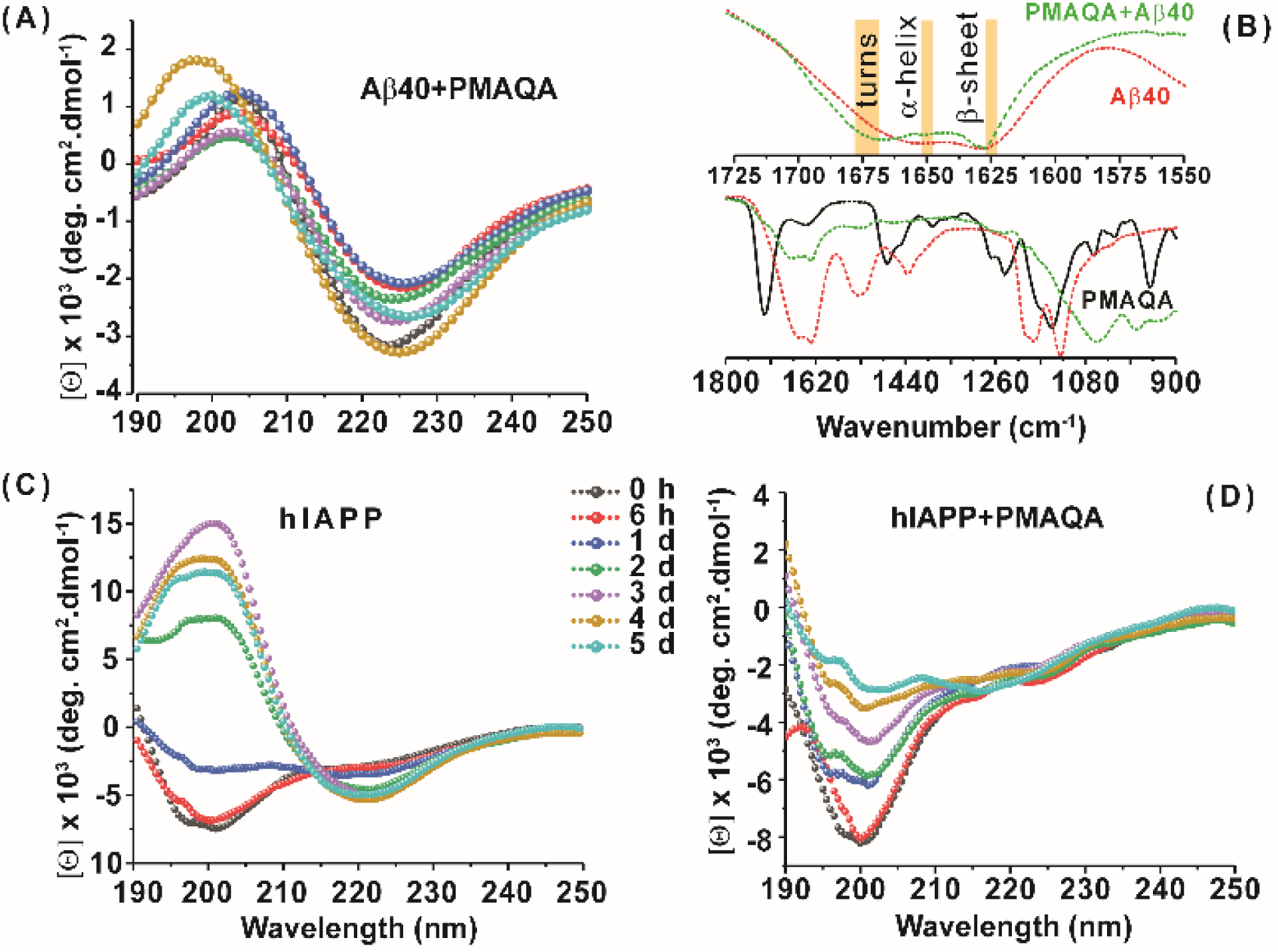
Time-lapse secondary structure change in Aβ40 and hIAPP. (A, C, D) Far-UV CD measurement monitoring the structural changes in 25 μM of Aβ40 (dissolved in 10 mM sodium phosphate buffer, pH 7.4) or hIAPP (30 mM sodium acetate buffer, pH 5.4) incubated with 37.5 μM PMAQA over 5 days. (B) FT-IR spectra (900-1800 cm-^1^) of Aβ40 in presence or absence of PMAQA incubated for 6 hours at room temperature. The peak shifting in the secondary structure (highlighted in yellow) fingerprint regions (1550-1725 cm-^1^) for Aβ40 upon PMAQA binding is shown on the top.

Similar to the amyloidogenic property of Aβ40, hIAPP in solution after 2 days depicted a CD spectrum maxima and minima at ≈200 and 218 nm, respectively, indicating its transitory state characterized by an increasing percentage of β-sheet (54%) (Fig. 2C, Table 1). But, unlike Aβ40, and as observed in ThT assays (Fig. 1D), CD spectra showed a relatively slow change in hIAPP’s secondary structure with a major CD minimum centered ≈200 nm up to day 3 (Fig. 2D). Secondary structure assessment of PMAQA bound hIAPP from CD spectra showed a relative increase in α-helix (14.5%) and decrease in β-sheet (21.7%) as compared to that observed in absence of PMAQA (Table 1). In addition, an increased percentage of parallel β-sheets was observed for hIAPP in presence of PMAQA over time (1-5 days) (13.4 % as compared to 0% in solution). This observation indicates that PMAQA bound hIAPP could have both anti-parallel and parallel β-structures. Such observations have been found previously using X-ray crystallography in different segments of hIAPP (segment 13-18 with parallel and anti-parallel β-structures for segments 16-21, 22-28, and 23-29).^30^ On the other hand, Aβ40 showing a positive and negative CD bands at ≈200 and 225 nm, respectively, on day 5 (transition from 222 nm observed on day 1) in presence of PMAQA indicates the formation of a predominant β-sheet structure (Fig. 2A). The observed Far-CD minimum with a red-shift of CD minimum (≈225 nm) in Aβ40 correlates to previously observed β-sheet rich supramolecular structures in modified Aβ and other small peptide aggregates.^31,32^

**Table 1.** Secondary structure assessment (%) of hIAPP from CD spectra in absence or presence of PMAQA by BESTSEL.^27^

| Time | 0 d | 1 d | 2 d | 3 d | 4 d | 5 d |
| --- | --- | --- | --- | --- | --- | --- |
| ***hIAPP in 30 mM sodium acetate pH=5.4*** |  |  |  |  |  |  |
| α-helix | 0.7 | 1.4 | 0 | 0 | 1.3 | 0.4 |
| $\beta^{\zeta}$ | 42.1 | 37.5 | 54.0 | 59.0 | 53.1 | 53.0 |
| $\beta^{\psi}$ | 0 | 3.9 | 0 | 0 | 0.2 | 0 |
| ***hIAPP+PMAQA in 30 mM sodium acetate pH=5.4*** |  |  |  |  |  |  |
| α-helix | 14.5 | 9.1 | 10.6 | 5.3 | 5.3 | 6.6 |
| $\beta^{\zeta}$ | 21.7 | 29.7 | 28.1 | 30.3 | 33.4 | 25.4 |
| $\beta^{\psi}$ | 0 | 0 | 0 | 5.7 | 3.7 | 13.4 |

$\zeta$ : Antiparallel β-sheet; $\psi$ : Parallel β-sheet

FT-IR spectra of PMAQA mixed with Aβ40 or hIAPP further supported the CD observation. As shown in Fig. 2B, predominant β-sheet structures (with an increasing percentage of turns, band at 1675 cm-^1^) of Aβ40 were observed with a sharp amide I peak at 1628 cm-^1^. In contrast, FT-IR spectrum of hIAPP remained unchanged with a band at 1650 cm^-1^that corresponds to α-helical structure (Fig. S2B). Size-exclusion chromatography (SEC) analysis of PMAQA mixed with Aβ40 incubated for ∼5 minutes at room temperature at 1:1.5 molar ratio showed two different elution profiles. The fractions collected at ∼5 to 12 mL corresponds to amyloid fibers or protofibers and that collected at ∼20 to 25 mL are free polymers or low order or monomeric Aβ40^33^ (Fig. S3A). Remarkably, SEC profile for PMAQA-hIAPP mixed solution incubated over night at room temperature under gentle shaking presented lower order or monomeric hIAPP or free polymers eluted at ∼20 to 25 mL^34^ (Fig. S3B). Taken together, the above described experimental results presented a counter active role of PMAQA on Aβ40 and hIAPP aggregation.

Next, we studied the binding mechanism of PMAQA with Aβ40 or hIAPP using an integrated NMR and molecular dynamics (MD) simulation approach. ^1^ H NMR spectra of peptide amide region (H-N) in the absence of PMAQA showed a monomer or lower order aggregate state of Aβ40 or hIAPP characterized by the observation of a number of dispersed NMR peaks (Fig. 3A). A substantial change in ^1^ H NMR was identified when 1.2 µM PMAQA was titrated with 60 µM of

**Fig 3.**
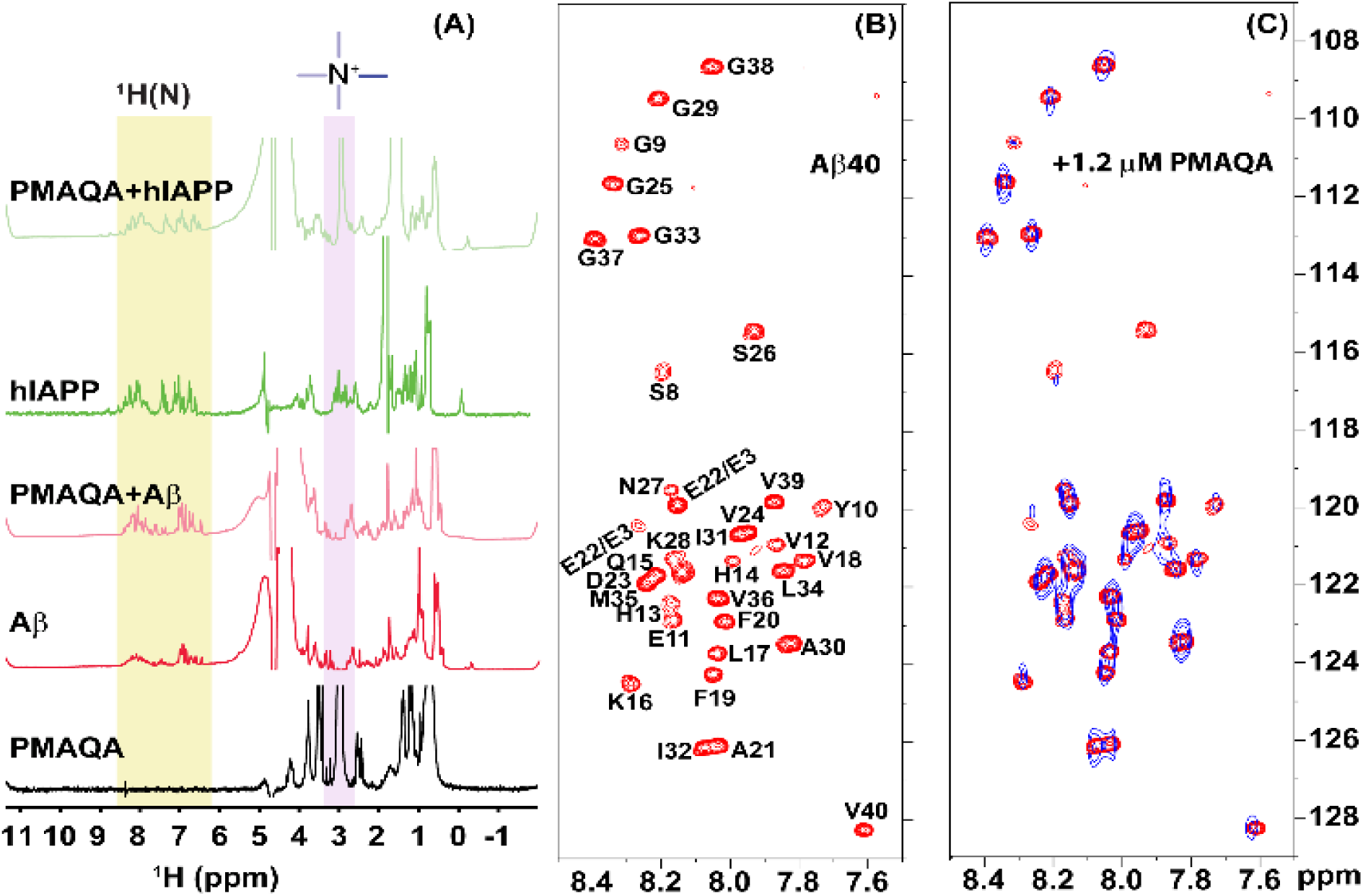
Interaction of Aβ40 or hIAPP with PMAQA studied using NMR. (A) 1D ^1^ H NMR spectra showing the interaction of PMAQA with Aβ40 (60 µM Aβ40, 1.2 µM PMAQA) or hIAPP (60 µM hIAPP, 50 µM PMAQA). The change in the NMR signal intensity of the –NR_3+_ proton in PMAQA or protein amide region (H-N) is highlighted. 2D ^15^ N/^1^ H SOFAST-HMQC spectra of Aβ40 (60 μM) dissolved in 10 mM sodium phosphate buffer, pH 7.4, containing 10% D_2_O at 10 °C recorded at 600 MHz in (B) absence of PMAQA (red) or (C) in presence of 1.2 µM PMAQA (blue). Spectra with an increasing concentration of PAMQA are provided in the supporting information (Figure S4).

Remarkably, the loss of amide proton peaks of Aβ40 by slightly increasing PMAQA concentration (to 3 µM) indicated an aggregation of Aβ40 (Fig. S4). In contrast, the amide-NH peaks of hIAPP (60 µM) were observed even when titrated with as high as 50 µM PMAQA. These NMR findings are in agreement with the observed conformational transition from CD and ThT based aggregation results (Figs. 1 and 2). Interestingly, the ^1^ H NMR observations revealed a significant line broadening for the proton peak of PMAQA’s –NR_3_ ^+^ at 2.97 ppm in Aβ40 solution indicating its interaction with peptide (Fig. 3A, light red). In contrast, a sharp proton peak for – NR_3_ ^+^ was observed in hIAPP solution (Fig. 3A, light green). This observation most likely indicates the formation of an electrostatic bond between the cationic PMAQA and Aβ40, as Aβ40 (at pH=7.4) and hIAPP (at pH=5.4) carry negative (∼ -2.8) and positive (∼ +2.9) charges, respectively. Thus, while the cationic group of PMAQA binds strongly to anionic Aβ40, a repulsive force could be expected in presence of cationic hIAPP.

To gain further insight into the mechanism of Aβ40 aggregation upon PAMQA binding, we performed 2D ^15^ N/^1^ H SOFAST-HMQC NMR titrations to map the polymer binding sites on Aβ40. As shown in Fig. 3B-C, a reduction of NMR signal intensities and change in chemical shifts of Aβ40 residues were observed indicating a PMAQA induced structural rearrangement for Aβ40. Residues mapping showed a substantial loss of Glu22/Glu3, Ser8 and Ser26 NMR signal intensities indicating a potential site of PMAQA’s interaction with Aβ40 (Fig. S4). Remarkably, an increase in the concentration of PMAQA resulted in a substantial line broadening, and loss of ^15^ N/^1^ H resonances of Aβ40 were observed at 3 µM PMAQA (Fig. S3).

To further explore the binding mechanism of PMAQA with hIAPP or Aβ40 at atomic level, we performed all-atom MD simulation on a time scale of 0.7 or 1 µs, respectively. Structural analysis showed a substantial number of hydrogen bond formation between PMAQA and Aβ40 over a time-scale of 1 µs (Fig. S5A and B). All-atom MD simulation revealed potential hydrogen bond and electrostatic interactions between the Glu22 or Asp23 residue (Aβ40) and polymer’s –NR_3_ ^+^ (Fig. 4A), whereas no interaction with Aβ40’s charged residues in the N-terminal (Asp1, Glu3 and Asp7) were observed. This is consistent with NMR results that showed a very little resonance change for the N-terminal residues (Fig. 3C). On the other hand, other residues such as Leu17, Phe20, Val24, Ser26, Asn27, Ile31, and Ile32 were observed to interact with PMAQA through hydrogen bond or hydrophobic interaction (Fig 4A, Table S1). ^1^ H/^15^ N NMR signal intensities measured from SOFAST-HMQC spectra showed a significant reduction in signal intensity for K16-D23 and A30-V40 regions (Fig. 3C). These observations are also in line with MD simulation based interaction map that indicated several intermolecular hydrogen bonds between Aβ40 and PMAQA (Table S1, Fig. S5A). Similarly, the MD calculations revealed several intermolecular hydrogen bond and hydrophobic interactions between hIAPP and PMAQA. hIAPP residues such as Asn21, Ile26, Ser28, Thr30, Asn31, Ser34 and Tyr37 were identified to be involved in intermolecular hydrogen bonding interactions with PMAQA (Fig. 4B, Table S2). Overall, PMAQA exhibited more number of hydrogen bonds with hIAPP than with Aβ40 indicating a relatively stronger binding affinity of PMAQA to hIAPP (Fig. S5A).

**Fig 4.**
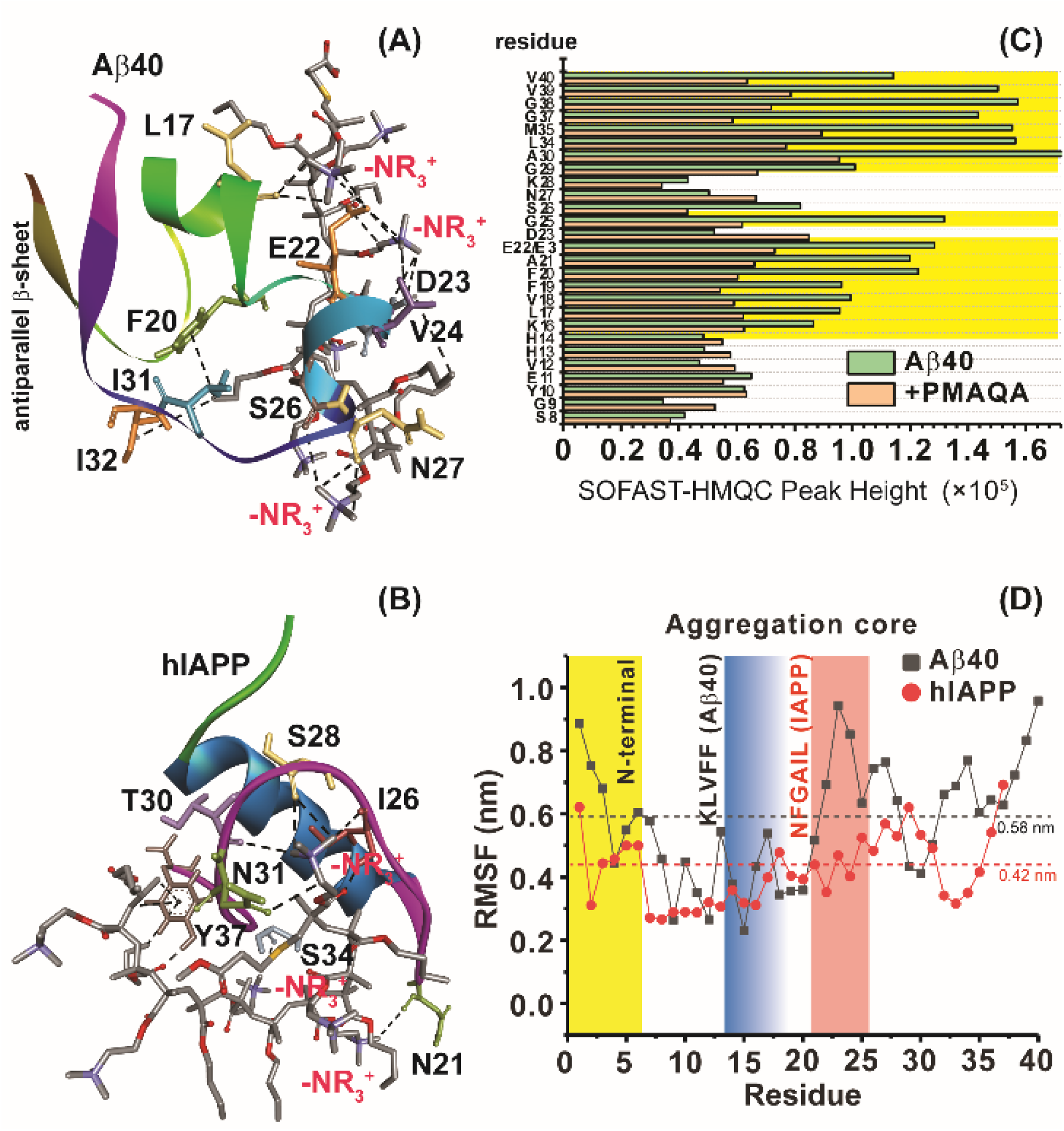
Structural insights into the PMAQA interaction with Aβ40 or hIAPP. MD snapshots showing PMAQA (shown in ball and sticks) interaction with Aβ40 (A) or hIAPP (B) shown as cartoon. The PMAQA binding peptide amino acids (shown in sticks) are labeled and hydrogen bonds are shown in black dashed lines in Discovery Studio Visualizer. (C) Signal intensities measured from SOFAST-HMQC spectra (see Figure 3B) of 60 µM Aβ40 and 0.6 µM PMAQA. The yellow area highlights the Aβ40‘s residues with significantly reduced signal intensities. (D) Root mean square fluctuation (RMSF) of residues in Aβ40 (grey) or hIAPP (red) interacting with PMAQA derived from 1 or 0.7 µs MD simulations, respectively. The yellow region indicates a comparatively flexible Aβ40 N-terminal domain. The blue and orange regions indicate aggregation core domains of hIAPP and Aβ40, respectively. The average RMSF value is shown using dashed horizontal lines.

The root mean square deviation (RMSD) of the backbone atoms calculated from 1 µs MD simulation of Aβ40-PMAQA system showed an RMSD plateau with an average value ≈11 Å (Fig. S5B). A relatively small backbone RMSD (average value ≈8.8 Å) was observed for hIAPP-PMAQA system indicating a comparatively stable complex formation (Fig. S5C). The RMSD of PMAQA calculated from all-atoms depicted a nearly equal RMSD value ≈6.5 Å for both MD systems (Fig. S5D). Overall, the stable protein backbone and PMAQA RMSD indicate a strong coupling with hIAPP or Aβ40 over the ∼µs time scale. Further the root mean square fluctuation (RMSF) analysis of individual amino acids in Aβ40 and hIAPP highlighted the potential PMAQA interaction regions. As illustrated in Fig. 4D, the N-terminal residues in Aβ40 were observed to be more flexible as compared to hIAPP when bound to PMAQA. The amyloid aggregation core domains in both Aβ40 (KLVFF) and hIAPP (NFGAIL) depicted an RMSF value lower than their corresponding average values as indicated in Fig. 4D. In hIAPP, residues 7-16 showed the lowest RMSF values that folded to a stable α-helical conformation. This indicates the PMAQA interaction restrain the structural and dynamic properties of hIAPP. The tight coupling of PMAQA close to the proximity of “NFGAIL” domain in hIAPP followed by restriction in protein structural rearrangement reveal mechanistic insights into PMAQA’s antagonist property in hIAPP aggregation. Further, the secondary structure analysis evolved from MD trajectories in hIAPP-PMAQA complex showed no significant secondary structural change during the MD simulation in hIAPP (Fig. S6, bottom). On the other hand, a substantial secondary structure change was observed in Aβ40 complexed with PMAQA including the induction of an anti-parallel β-sheet along the terminal residues (Figs. 4A and S6, top). The interaction of PMAQA with the centrally located residues of Aβ40 such as Glu22, Asp23 and Val24 induced a random-coil to β-structure transition in Aβ40. Structural changes in Aβ by affecting these central residues by mutation (E22G), disruption of the D23-K28 salt-bridge or binding of hexapeptide of genetic Aβ variants has been reported previously to be crucial in promoting Aβ40 aggregation and induction of terminal β-structure.^35–38^

In conclusion, we have demonstrated the counter activities of a cationic PMAQA polymer on two different amyloidogenic peptides that are connected to AD and T2D. At sub-micromolar concentration, PMAQA showed significant inhibitory activity in hIAPP aggregation; whereas it significantly accelerated Aβ40’s aggregation by quickly altering the equilibrium state of Aβ40 from an unfolded structure to a β-sheet structure. Our mechanistic study provides insights into the binding of PMAQA to Aβ40 or hIAPP at atomic-level that could be helpful in understanding the modulation of peptide self-assembly and could aid in potential inhibitor designing. We believe that the opposite aggregation kinetics of two different amyloidogenic proteins in the presence of a cationic polymer delineated in this study likely to open avenues to test their potential therapeutic activities against an array of amyloid proteins involved in other human amyloid diseases by controlling functionalization of the polymer’s chemical property.

## Methods

### Materials

The polymethacrylate quaternary ammonium copolymer (PMAQA, ∼4.7 kDa) was synthesized and purified as reported elsewhere.^24^ Unlabeled and uniform ^15^ N isotope labeled full-length Aβ40 (DAEFRHDSGYEVHHQKLVFFAEDVGSNKGAIIGLMVGGVV) was recombinantly expressed in *E. coli* BL21 (DE3). The Aβ40 expression was followed from previously described protocols ^39,40^ and purified by loading the samples to an ^ECO^PLUS HPLC column packed with the reversed-phase separation material. Synthetic hIAPP (KCNTATCATQRLANFLVHSSN NFGAILSSTNVGSNTY-NH2) was purchased from AnaSpec at > 95% purity.

### Sample preparation

The Aβ40 peptides were dissolved in 5% (v/v) NH_4_OH and lyophilized at a concentration of 0.1 mg/ml. The Aβ40 peptide powder was re-suspended in 10 mM sodium phosphate, pH 7.4 and sonicated for 30s followed by centrifugation at 14,000 × *g* for 15 min at 4 °C to remove small aggregates. 1 mg/mL hIAPP was treated with 1,1,1,3,3,3-hexafluoroisopropanol (HFIP) and kept on ice for 30 minutes. The peptide solutions were aliquoted to 0.1 mg/mL and lyophilized. The hIAPP powder was re-suspended in 30 mM sodium acetate buffer, pH 5.4 and sonicated for 30s at 4 °C. The PMAQA powder (10 mg/mL) was dissolved in deionized water.

### Circular dichroism and Fourier transform-infrared spectroscopy

Aβ40 or hIAPP secondary structural transition in presence and absence of PMAQA was studied by Far-UV circular dichroism (CD) using a JASCO (J820) spectropolarimeter. A light-path length (1 mm) cuvette containing 25 μM peptide (Aβ40 or hIAPP) solution titrated with an increasing concentration of PMAQA (0.25 to 50 μM) was used to monitor the evolution of structural transition at 25 °C. The samples were stored at room temperature and the Far-CD spectra were recorded for 5 days at different time intervals. The CD spectra were averaged and expressed as the mean residue ellipticity [Θ] after subtracting the signal from a solution without peptide.

Fourier transform-infrared (FT-IR) spectra were measured for Aβ40 or hIAPP mixed with PMAQA (incubated for 6 hours) at 1:1 molar ratio in transmission mode within a range of 4000– 400 cm^-1^ using a Thermos scientific ATR-FTIR instrument. Samples were lyophilized for 72h to remove water followed by FT-IR measurement.

### Thioflavin-T fluorescence assay

Thioflavin T (ThT) fluorescence assays were performed to monitor the aggregation kinetics of 5 μM Aβ40 or hIAPP at 37 °C in presence of PMAQA (1, 2.5, 5, 10, and 50 μM) and 10 μM ThT. Fisher 96-well polystyrene plates with a sample volume of 100 μl/well were used for the ThT assay. The kinetics of amyloid formation was monitored at 3-min intervals with no-shaking conditions for 4 days using a microplate reader (Biotek Synergy 2) with an excitation and emission wavelengths of 440 and 485 nm, respectively. The aggregation kinetics was interpreted by fitting the ThT curves using the following equation.^41^

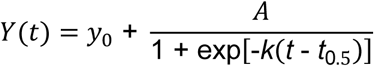

Where *y*_0_is the pre-transition baseline, *k* is the apparent growth rate constant and *t_0.5_* is the half-time when ThT fluorescence reaches half of its maximum intensity. The lag-time (t_lag_) is defined as t_lag_=*t*_0.5_-(1/2*k*).

### SEC

The size profiling of Aβ40 or hIAPP species bound to PMAQA were studied by passing them through SEC using a Superdex 200 Increase 10/300 GL column operated on an AKTA purifier (GE Healthcare, Freiburg, Germany). 10 µM of freshly dissolved Aβ40 was mixed with 15 µM of PMAQA and incubated for 5 minutes at room temperature before loading to SEC column. Simllarly, 10 µM of freshly dissolved hIAPP was mixed with 10 µM of PMAQA and incubated over night at room temperature with gentle shaking at 300 rpm before loading to SEC column.

### NMR

1D and 2D NMR spectra were recorded on a 600 MHz Bruker Avance III NMR spectrometer equipped with a z-axis gradient cryogenic probe. Unlabeled hIAPP, Aβ40 or ^15^ N-labeled Aβ40 peptides (peptide concentration = 60 μM) dissolved in 10 mM sodium phosphate, pH 7.4 (for Aβ40) or 30 mM sodium acetate, pH 5.4 (for hIAPP) buffer containing 90% H_2_O/10% ^2^ H_2_O was used for NMR measurements. The 2D ^15^ N/^1^ H SOFAST-HMQC (Bruker *sfhmqcf3gpph* pulse program)^42^ NMR titration experiments of Aβ40 (60 μM) with 0.6, 1.2 and 3 μM PMAQA were recorded at 10 °C with 64 scans and 200 t1 increments. The NMR spectra were processed using TopSpin 3.5 (Bruker) and analyzed using Sparky.^43^

### MD simulations

The 2D structure of PMAQA was created using ChemDraw 16.0 and were exported to Chem3D for energy minimization followed by molecular dynamics (MD) calculations using MMFF94 force field.^44^ The 3D structure of PMAQA and its topology files were created using ATB builder^45^ for all-atom MD simulation. The solution NMR structures of Aβ40^46^ (PDB ID: 2LFM) and hIAPP^47^ (PDB ID: 5MGQ) were considered as the initial structure for PMAQA interaction analysis. The MD system was built in GROMACS^48^ software package, version 5.0.7 (GROMOS96 54A7^49^ force field), by placing Aβ40 or hIAPP at the center of a cubic box and the polymer ∼ 1 nm away from the protein. The MD systems were solvated using SPC/E water (≈ 1000 kg m^-3^) and neutralized by adding counter ions followed by energy-minimization using the steepest-descent method. Short NVT (100 ps) followed by 1 ns NPT (310 K and 1 bar) was performed to equilibrate the MD systems. MD simulations were carried out using 3D periodic boundary conditions over a production run of 0.7 and 1µs for hAIPP-PMAQA and Aβ40-PMAQA systems, respectively. MD trajectories were interpreted using visual molecular dynamics^50^ and images were built using Discovery studio visualizer 3.5 ^51^.

## Acknowledgements

This study was supported by funds from NIH (AG048934 to A.R.). This work was (in part) performed under the International Collaborative Research Program of Institute for Protein Research, Osaka University, ICR-18-02. We thank Professor Toshimichi Fujiwara in the Institute for Protein Research, Osaka University, for providing parallel computing facility on SGI UV 3000. We thank Professor Bernd Reif for providing us the recombinant expression system and protocol for the production of amyloid-beta (1-40) peptide.

